# Silk assembly integrates cells into a 3D fibrillar network that promotes cell spreading and proliferation

**DOI:** 10.1101/403345

**Authors:** Ulrika Johansson, Mona Widhe, Nancy Dekki Shalaly, Irene Linares Arregui, Linnea Nilebäck, Christos Panagiotis Tasiopoulos, Carolina Åstrand, Per-Olof Berggren, Christian Gasser, My Hedhammar

## Abstract

Tissues are built of cells integrated in an extracellular matrix (ECM) which provides a three-dimensional (3D) fibrillar network with specific sites for cell anchorage. By genetic engineering, motifs from the ECM can be functionally fused to recombinant silk proteins. Such a silk protein, FN-silk, which harbours a motif from fibronectin, has the ability to self-assemble into fibrillar networks under physiological-like conditions. Herein we describe a method by which mammalian cells are added to the silk solution before assembly, and thereby get uniformly integrated between the formed fibrils. In the resulting 3D scaffold, the cells proliferate and spread out with tissue-like morphology. Elongated cells containing filamentous actin and defined focal adhesion points confirm proper cell attachment to the FN-silk. The cells remain viable in culture for at least 90 days. The method is also scalable to macro-sized 3D cultures. Silk fibers with integrated cells are both strong and extendable, with mechanical properties similar to that of artery walls. The described method enables both differentiation of stem- or precursor cells in 3D and facile co-culture of several different cell types. We show that inclusion of endothelial cells leads to the formation of vessel-like structures throughout the tissue constructs. Hence, silk-assembly in presence of cells constitutes a viable option for 3D culture of cells integrated in a fibrillary ECM-like network, with potential as base for engineering of functional tissue.

---

Since the 1940s, *in vitro* cultures of mammalian cells have become indispensable for both basic research and industrial applications. Most cell culture studies are today performed on hard plastic or glass surfaces because of the ease, convenience and high viability associated with this method. However, forcing cells to adapt against a flat and rigid 2D surface means that almost half of their surface area is dedicated to adhesion, whereas cells in the body are likely to receive other signals not just at their ventral surface but in all three dimensions. This can alter the cell metabolism and functionality, thereby providing results different from what would be obtained from cells in their natural *in vivo* environment [1]. Lately, the bearing of culturing cells in 3D has been increasingly acknowledged, and it is expected that 3D cultures provides cellular responses that are of higher biological relevance. When comparing cells cultured in 2D versus 3D, significant differences associated with key biological processes such as adhesion, proliferation, differentiation *etc.* have been found [2-5].

Promoting cells to grow and form 3D structures *in vitro* has been proven more difficult than first anticipated. By forcing cell-cell contacts to form using *e.g.* non-adhesive surfaces and gravity, cells can be gathered into spheroids [6]. However, spheroids are typically irregular and limited in size, since the cells lack a supporting matrix. To permit culture of larger 3D cell arrangements, a temporary support is needed, and for this a range of scaffolds made of both natural and synthetic materials have been developed [7]. Traditionally, a top-down approach has been used, where cells are seeded on top of pre-made scaffolds. Although the scaffolds thereby can be constructed to provide sufficient mechanical support, this approach has limitations in low seeding efficiency and non-uniform cell distribution leading to restricted cell-cell contacts [8]. As alternative, a bottom-up approach has been initiated, relying on assembly from soluble components together with the cells [9]. The choice of material is then limited to those which can be assembled under conditions compatible with cell viability. Several strategies for creation of cellular constructs, *e.g.* rapid prototyping and solid free form fabrication, have been developed during the last years [10, 11]. One exciting route is 3D printing of hydrogel-based bioinks containing cells [12]. For this, hydrogels which can be induced by mild conditions, *e.g.* addition of calcium ions [13], are used. Thereby the cells are encapsulated in a 3D environment that is beneficial in terms of a high water content. However, from the perspective of a cell, the microenvironment in a pliable reticular hydrogel is very different compared to the native ECM. *In vivo,* the cells are surrounded by fibrillar networks of ECM to which they can firmly attach and transmit mechanical forces through [14]. Despite the possibility to fine tune the bulk mechanical properties of many hydrogels, such polymer chains are too thin to allow proper establishment of cell-matrix interactions.

We have previously developed a scalable process for recombinant production of the spider silk protein 4RepCT [15, 16]. This protein has the unique ability to self-assemble into biocompatible fibrillar networks in aqueous physiological-like buffers at room temperature [16, 17]. Further, 4RepCT has been functionalized at the genetic level with a cell adhesion motif from fibronectin (FN) to allow formation of FN-silk, that promotes firm cell attachment [18]. Previous studies of cells seeded on already formed silk scaffolds showed good proliferation and migration, although cells then grew along the fibrillar surfaces of the silk rather than inside [18, 19]. A proper balance and distribution of cells within the scaffold is crucial for formation of cultures with potential to have tissue-like properties. Our recent discovery about how the air-water interface can be used to promote silk assembly [20] has opened up for a new possibility to efficiently integrate cells into a material. Herein we explore the option to include cells already during assembly of FN-silk, with the scope to develop a method for 3D culture of cells integrated in a fibrillar network.

## Silk-assembly allows for direct integration of cells into 3D fibrillar networks

In solution, the recombinant silk protein 4RepCT forms dimers of mainly helical/random coil structure [21]. In order to promote a conformational rearrangement driving assembly into a multimeric silk state, alignment against an interface, for example air-water, can be used [20]. The easiest way to formulate silk is thus accomplished by gentle introduction of air bubbles into the solution of silk protein, giving rise to a 3D foam with randomly distributed fibrillar walls [19]. To examine the feasibility of this method for formation of a silk foam with integrated cells (Fig. 1a, I-III), we first added a drop of dispersed cells (mouse mesenchymal stem cells) to the FN-silk protein solution before assembly. Upon gentle introduction of air bubbles, an expanding wet foam structure was obtained, with similar appearance as a foam from the silk protein solution by itself. After 30 minutes in the cell incubator, the formed foam had settled and remained stable as a cell-containing 3D scaffold even when covered with culture media (Fig. 1a, IV). Throughout the following culture period, the foam became denser and less transparent, as trapped air bubbles were released and the integrated cells increased in number (Fig.1a, V). Already after 3 days in culture, cells were found spread throughout all dimensions of the foam (Fig. 1a, VI).

**Figure 1.**
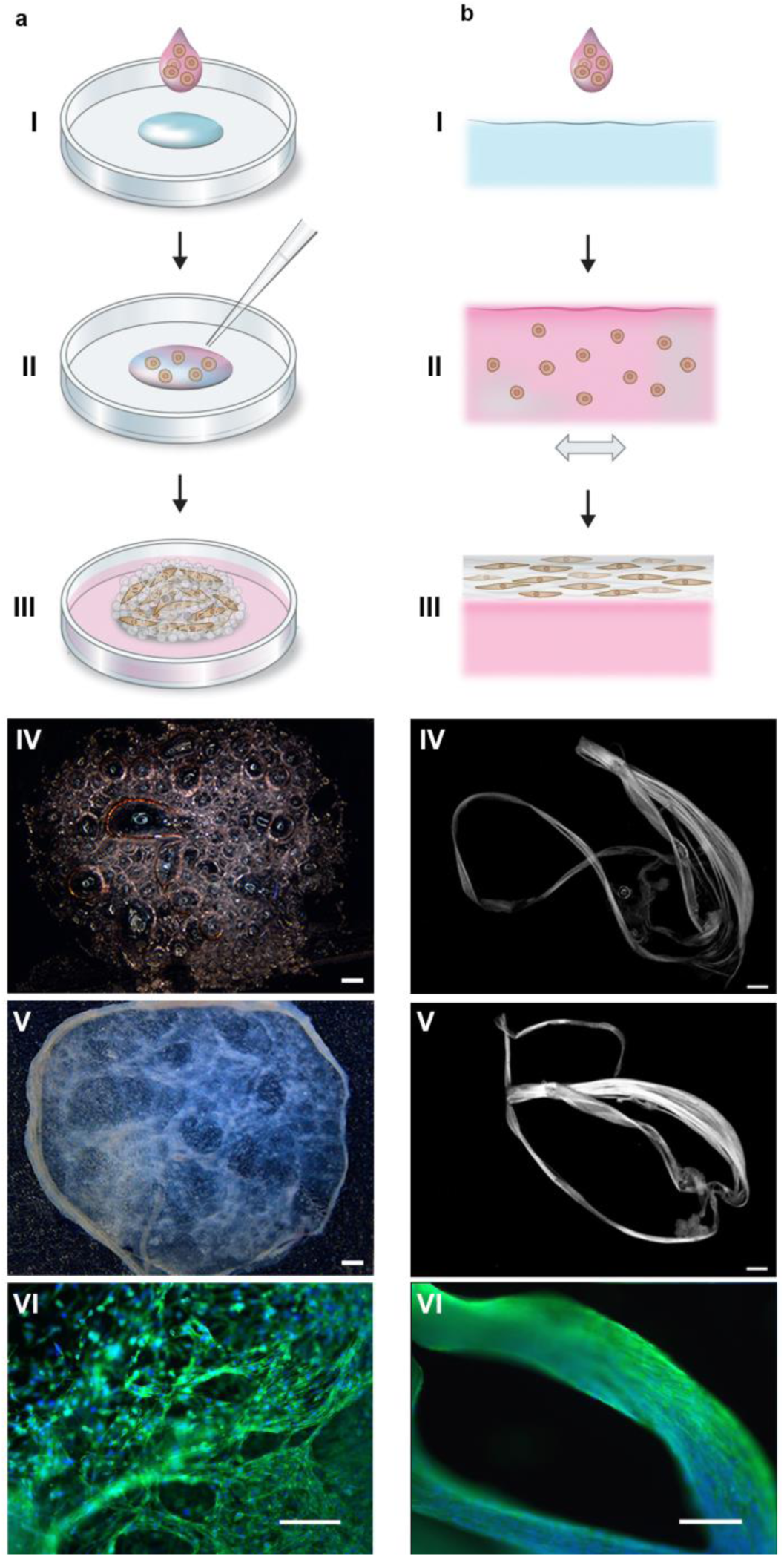
Silk-assembly to integrate cells into 3D fibrillar scaffolds. **(a)** Schematic description of the formulation of silk foam with integrated cells. Cells suspended in culture medium (pink) are added to a drop of FN-silk protein solution (blue) placed in the middle of a non-treated culture well (I). Air bubbles are gently introduced through a pipette tip (II), to give rise to a 3D foam with cells. After 30 minutes in the cell incubator, additional culture medium is added to cover the foam completely (III). Day 1 after formulation, the silk foam with cells looks almost transparent, although harboring air bubbles (disappear after media change) (IV). After 2 weeks of culture, the foam with integrated cells has stabilized and shows a denser appearance (V). Already at day 3 the foam is filled with well-spread cells (here mouse mesenchymal stem cells (MMSC)) (VI). Actin filaments are visualized by phalloidin (green) and cell nuclei by Dapi staining (blue). Scale bar IV-V =1 mm, VI = 100 μm. **(b)** Schematic description of formulation of silk fibers with integrated cells. Cells suspended in medium (pink) are added to the FN-silk protein solution (blue) (I). During gentle repeated uniaxial tilting for 1-3 hours (II) the silk proteins assemble at the air-liquid interface into macroscopic fiber bundles with incorporated cells (III). The silk fibers with cells are easily retrieved (IV) and can be placed in a well for further culture, whereby the thickness increases over 2 weeks (V). At day 3, aligned cells (here MMSC) are found spread integrated in the fiber bundle (VI). Actin filaments are visualized by phalloidin (green) and cell nuclei by Dapi staining (blue). Scale bar IV-V = 1 mm, VI = 100 μm.

The possibility of directing the alignment of cells has been recognized as an essential component of biomimicry [11]. In the foam formulation process, the silk-assembly triggered by the air bubbles results in a 3D network of fibrils. This network resembles random structures formed in nature, such as the sparsely organized networks of ECM found in loose connective tissue [22]. However, hierarchically ordered structures, with alignment of cells into a certain orientation, is necessary for reconstruction of line-shaped tissues such as blood vessels, nerves, muscle fibers and tendons [8, 23]. A gentle repeated uniaxial tilting of the silk protein solution during the assembly process gives rise to aligned silk structures in the format of bundles of fibrils formed at the air-water interface [15]. Encouraged by the cell compatibility of the foam formulation method, we added cells (mouse mesenchymal stem cells) also to the FN-silk solution before inducing fiber formation by repeated tilting of the interface (Fig. 1b, I-III). Macroscopic fibers started to become visible within the same time frame as seen for silk protein alone (∼30 minutes) [24] and continued to grow the following hours. To the naked eye, the formed fiber bundles with integrated cells first looked similar to those without cells, but continued to grow in thickness during the culture period (Fig. 1b, IV-V). Aligned cells, with prominent actin filaments, were found tightly integrated within the fiber bundle (Fig. 1b, VI).

As an alternative format, we investigated if high-density cell sheets could be gathered by combining the solution of FN-silk protein with a high concentration of cells, without promoting silk-assembly in any direction. By keeping a drop of FN-silk protein containing cells still in the middle of a non-treated culture plate for 30-60 minutes, silk-assembly at the interfaces had stabilized a sheet with integrated cells, which could be covered with culture media for further culture (Suppl. Fig. 1). A majority of the cells within the silk sheet remained viable.

In order to examine the versatility of the silk assembly method, 11 different cell types (see Suppl. Table 1) were investigated for incorporation during the silk formulation process. The studied cell types, mainly isolated human primary cells, were selected to include a wide range of adherent cells, originating from various tissues of the body. The cell to ECM ratio varies widely depending on the tissue [25] and in some tissues it is of importance that cells are growing densely and in contact with each other. With the silk-assembly method, the added amount of cells, and thus the density, can easily be adjusted to match ranges from the rather cell sparse cartilage to high-density tissues such as liver, with up to 10^8^-10^9^ cells/cm^3^ [26]. The very simple procedure for formation of a silk foam allows for a high cell seeding efficiency, since all the added cells are integrated by the silk assembly process. It is thus a suitable procedure also for 3D culture of cells available only in minute amounts.

## Cells are highly proliferative within 3D silk

The herein described process for formation of silk scaffolds with integrated cells utilizes silk assembly at the air-water interface, which does not require any additional reagents or conditions known to harm cells (*e.g.* changes in pH, temperature, ions and radiation). Still, the entrapment itself might affect the ability of cells to migrate and expand. Therefore, we performed thorough investigations of the growth profiles of the different cells types within both foam and fibers (Fig. 2a,b, Suppl. Fig. 2). In all trials performed, increasing metabolic activity of cells within the 3D silk could be observed over time, indicating cell proliferation. Typically, a faster expansion phase could be observed during the first days. Over time, the cell-containing silk scaffolds appeared denser, and microscopy analysis revealed that cells were distributed throughout the scaffolds (Suppl. Fig. 3).

**Figure 3.**
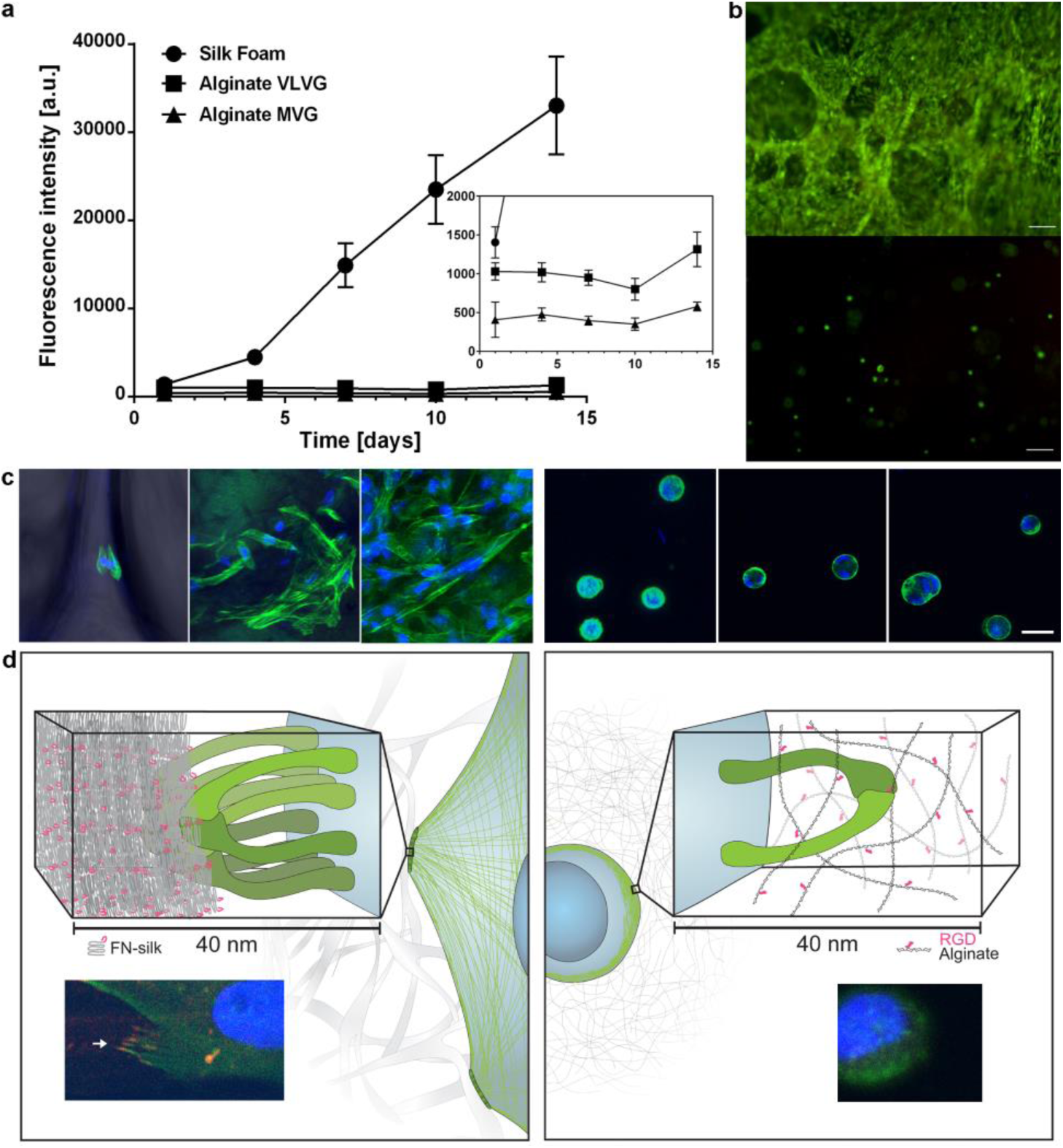
Attachment and expansion of cells within FN-silk compared to an RGD-coupled hydrogel. **(a)** Representative graph (mean and standard deviation) of Alamar Blue viability assay showing metabolic activity of fibroblasts (HDF) within FN-silk foam (circle), a low viscosity (VLVG) alginate hydrogel coupled with RGD (square), and a medium viscosity (MVG) alginate hydrogel coupled with RGD (triangle) during two weeks of culture. Insert shows a zoomed in view of the lower intensities. **(b)** Representative live (green) and dead (red) staining of human mesenchymal stem cells (HMSC) in FN-silk foam (upper) and RGD-coupled alginate VLVG (lower) at day 14. Scale bars = 100 μm. **(c)** Confocal scans of HMSCs integrated into FN-silk foam (left panel), and RGD coupled alginate hydrogel MVG (right panel) after 1h, 4 days and 7 days. Actin filaments are visualized by phalloidin staining (green) and cell nuclei are stained with Dapi (blue). Scale bars = 20 μm. **(d)** Schematic description of our hypothesis for the reason behind the observed difference in interactions between cells and silk (left) versus RGD-coupled alginate hydrogels (right). Several integrin pairs (green) can adhere and gather to the silk fibrils, forming focal adhesions at the edge of actin filaments, enabling the cell to spread and proliferate. In the alginate hydrogel, a single integrin pair (green) can bind to the coupled RGD-motif, but subsequent gathering into focal adhesions are restricted within the thin hydrogel network. Inserts show examples of a cell (fibroblast) after 3h in FN-silk foam (left) and a low viscosity (VLVG) alginate hydrogel coupled with RGD (right). Actin filaments are visualized by phalloidin staining (green), and focal adhesions can be seen where this is co-localized with staining for vinculin (red). Cell nuclei are stained with Dapi (blue).

**Figure 2.**
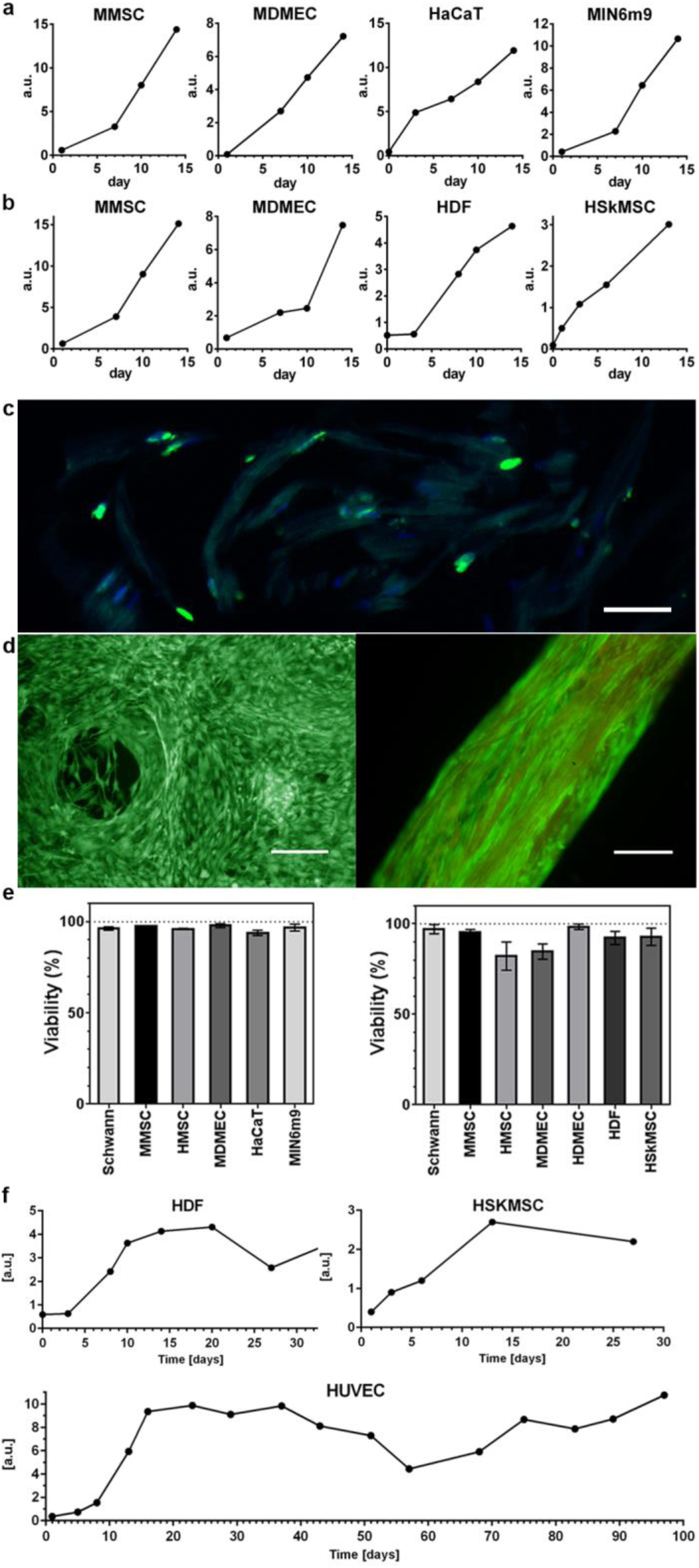
Proliferation and viability of cells integrated in 3D silk. Representative graphs of Alamar blue viability assay show increasing metabolic activity during the first 2 weeks within **(a)** foam (N=3-4, n=3-7), and **(b)** fibers (N=1-9, n=2-13), reflecting growth of the various integrated cell types (for cell abbreviations see Suppl. Table 1, for details, see Suppl. Table 2). **(c)** Cell division occurs deep within the 3D silk scaffolds. Cryosection of fiber with integrated fibroblasts (HDF) fixed at day 11 and stained with FITC-anti BrdU for newly synthesized DNA (green) and Dapi (blue). The silk shows a dim autofluorescence in the blue/green range. **(d)** Representative live (green) and dead (red) staining of mouse mesenchymal stem cells (MMSC) in foam (left) and HDF in fiber (right) at day 14. The fiber shows a dim autofluorescence in the red range. Scale bars = 100 μm. **(e)** Viability (%, mean and standard deviation) after 14 days culture of different cell types (see Suppl. Table 1) in foam (left graph), and in fibers (right graph) (N=1-3, n=4). **(f)** Long time cultures of cells integrated into fibers show maintained metabolic activity (Alamar blue) during the entire study period (up to 97 days)

To examine if also cells within the innermost part of the silk scaffolds were proliferating, the nucleotide analogue BrdU was added to allow incorporation into the genomes of dividing cells before fixation and cryosectioning. In this way, BrdU positive and thus proliferative cells present deep within the silk fibers could be observed at defined time points (Fig. 2c).

## Cell viability is high throughout the 3D silk

In order to constitute a workable option, it is important that the viability of the integrated cells is maintained during longer culture times. Therefore, a two-color fluorescence assay which simultaneously stains live (green) and dead (red) cells was applied to silk scaffolds assembled in presence of different cell types. Under the fluorescence microscope, it became evident that the majority of the cells were still alive after 2 weeks within the silk (Fig. d, Suppl. Fig. 4a). The overall viability at day 14, depending on cell type, was 94-98% in foam and 82-98% in fibers, respectively (Fig. 2e). Even after longer culture periods a majority of the cells were stained as viable (Suppl. Fig. 4b), which could be confirmed with the viability assay during longer culture periods (1-3 months) (Fig. 2f).

**Figure 4.**
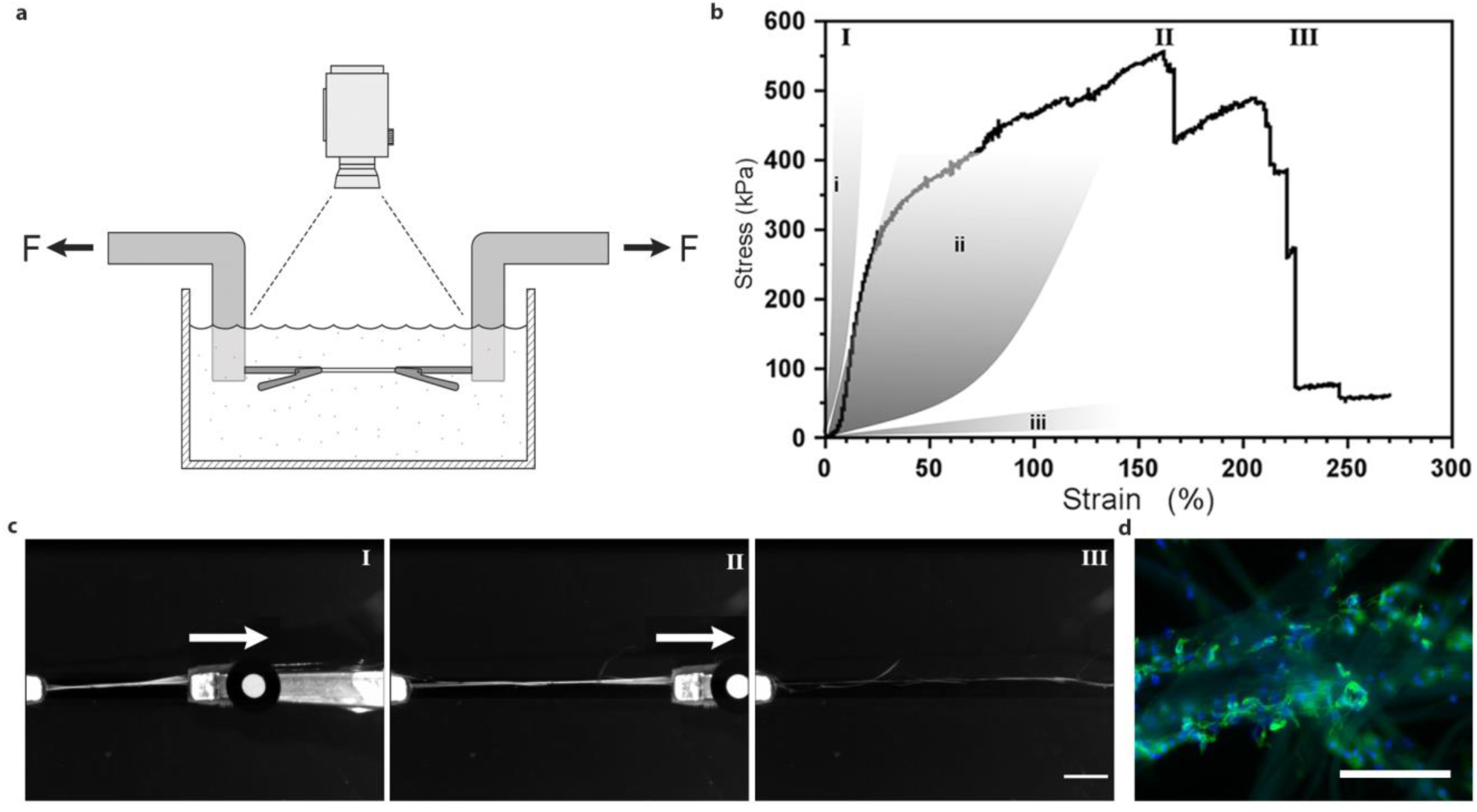
Uni-axial tensile testing of silk fibers with integrated mesenchymal stem cells. **(a)** Illustration of the experimental set-up for tensile tests performed in PBS buffer at 37°C in order to maintain viable cells. (**b**) Representative First Piola-Kirchhoff stress versus strain curve of a FN-silk fiber with integrated mesenchymal stem cells (MMSC) subjected to tensile testing after 14 days of culture. Stress-strain curve illustrates a rather linear (and probably elastic) phase that is followed by a plastic-like (irreversible) deformation phase until the maximum stress is reached, and the fiber breaks. For comparison, grey areas represent ranges of stress strain properties in tendons and ligaments (i), artery walls (ii), and brain tissue (iii). Roman numbers refer to images (**c**) taken during the tensile test, *i.e.* during start (I), extension (II) and breakage (III) of the fibers. Scale bar = 5 mm. (**d**) Micrographs of the breaking point of fibers with MMSCs after tensile testing. Actin filaments are visualized by phalloidin staining (green) and cell nuclei are stained with Dapi (blue). Scale bars = 200 μm.

## Comparison of cell spreading and proliferation within silk and alginate hydrogels

Encouraged by the positive growth profiles obtained from cells integrated into silk, we decided to perform parallel experiments with cells encapsulated in a hydrogel, for comparison. Alginate was chosen as a representative hydrogel since the gelation method is very mild and widely used for cell culture [13]. An alginate variant coupled with the cell binding motif RGD was chosen to provide cell adhesion sites comparable to the RGD-containing FN-motif. When performing parallel experiments with cells in FN-silk and alginate, respectively, the growth curves obtained were markedly different. A clear expansion phase was seen for cells integrated in silk, while cells encapsulated in alginate remained at an almost steady metabolic state (Fig. 3a, Suppl. Fig. 5a). This observation is in line with previous reports of limited proliferation of cells encapsulated in alginate hydrogels [27, 28]. Live/dead staining confirmed a high ratio of viable to dead cells 4 hours after seeding as well as after 2 weeks of culture in both scaffold types (Fig. 3b, Suppl. Fig. 5b).

**Figure 5.**
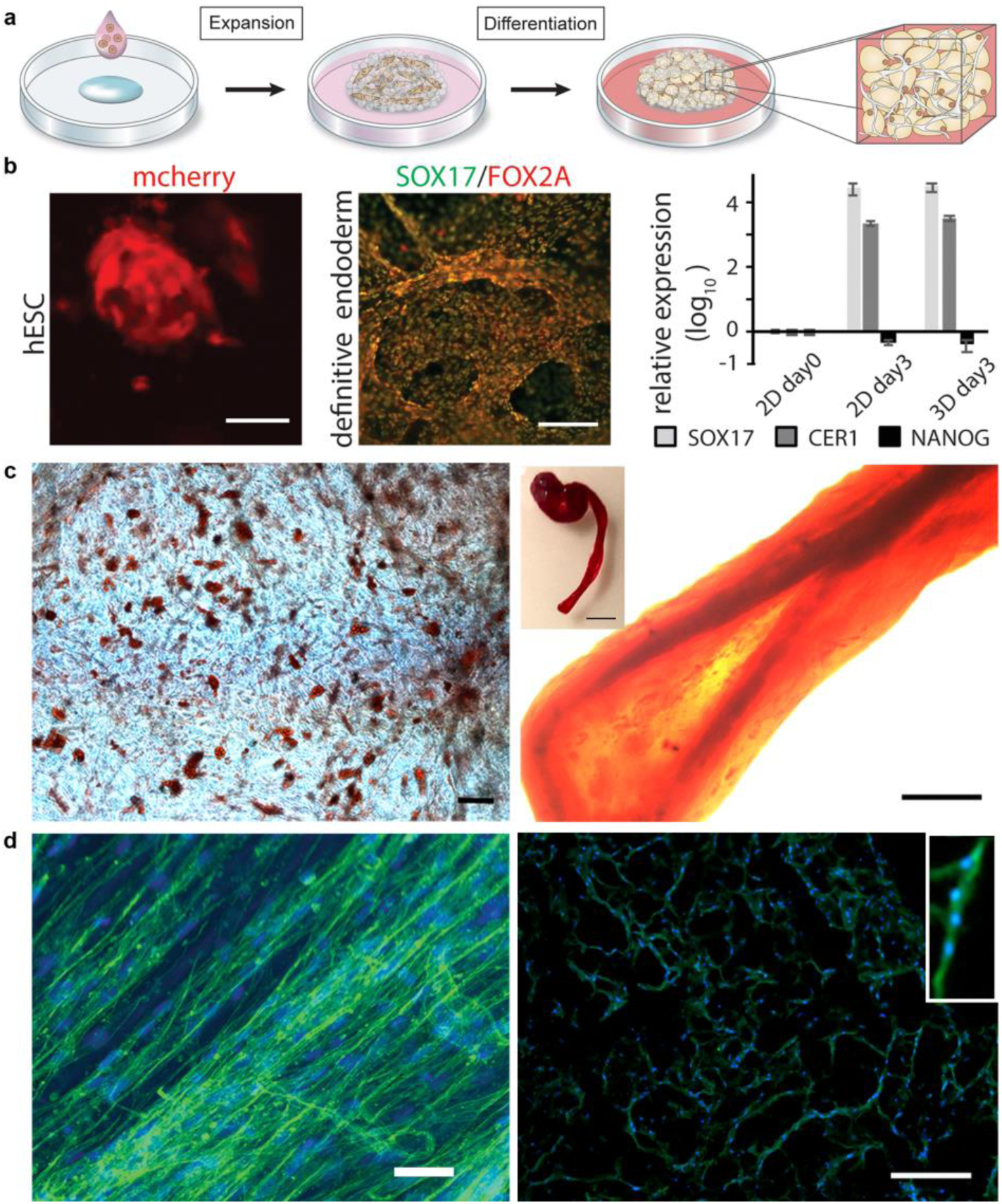
Differentiation of cells in 3D silk. **(a)** After initial expansion of stem cells integrated to 3D silk, differentiation into various tissue types can be triggered by addition of appropriate factors. **(b)** Differentiation of pluripotent stem cells. Left: Human embryonic stem cells (hESC) visualized by mCherry detection at 48h after cell integration into FN-silk foam. Scale bar = 50μm. Middle: Immunostaining for endodermal markers SOX17 (green) and FOX2A (red) after 3 days of differentiation. Scale bars = 200μm. Right: Gene expression (*SOX17*, *CER1,* NANOG) of hESC in a FN-silk foam compared to 2D culture, analyzed by RT-qPCR at day 3 of endodermal induction. Bars represent the mean fold change ±standard deviation (n=4). **(c)** Differentiation of multipotent adult stem cells. Left: Human mesenchymal stem cells (HMSC) in FN-silk foam differentiated into the adipogenic linage containing lipids, visualized by Red Oil staining (red) (N=2, n=4). Scale bar = 100 μm. Right: HMSCs differentiated into the osteogenic linage, probed with osteogenic marker for calcium content (Alizarin Red S (red) in FN-silk fiber (right, scale bar = 200 μm), (N=2, n=4). Inset shows photo of a whole fiber (right), scale bar = 1 mm). **(d)** Differentiation of adult precursor cells. Left: After 14 days in differentiation media, skeletal muscle satellite cells (HSkMSC) within a FN-silk fiber show prominent actin filaments, as visualized by phalloidin staining (green). Right: Myogenic differentiation of skeletal muscle satellite cells (HSkMSC) visualized by Desmin staining (green). Dapi-stained nuclei in blue. (N=9, n=4). Scale bars = 200 μm. A close up of the area of a multinucleated myotube is shown in the upper right corner.

In order to investigate the reason for the difference in growth behavior, we decided to look closer into the cell microenvironment at selected time points during the culture. Confocal microscopy analysis revealed that cells within silk (both foam and fiber) attained an elongated shape already after 1 hour (Fig. 3c, left, Suppl. Fig. 5c, left), indicating prompt attachment and spreading along the fibrils. When encapsulated within the hydrogel, the majority of the cells exhibited a rounded morphology throughout the culture period (Fig. 3c, right). After 4 and 7 days of culture, multiple nuclei could be observed within the rounded cell formations, indicating some proliferation but no spreading when encapsulated in the hydrogel. The cells integrated in silk were increasingly spread throughout the scaffolds during the culture period, with a clear directional alignment in the fibers (Suppl. Fig. 5c). An increase in cell number during culture in silk scaffolds could also be confirmed from the micrographs (Fig. 3c).

Based on the obtained results, we formulated a hypothesis to explain the requirements of the microenvironment to obtain tissue-like culture of cells in 3D (Fig. 3d). Hydrogels formed under mild conditions allow encapsulation of cells and can be used to obtain an even distribution of viable cells in 3D. The cells stay viable during culture, but with rounded cell morphology even though the alginate gel contains an extensive amount of cell binding motifs. This is likely due to the importance of appropriate dimensions and mechanical properties of the scaffold were the adhesion motif is presented, in order for the cells to attain an accurate cell morphology [29]. The cells need to assemble several integrins to form focal adhesions at the rear of their cytoskeletal structures to trigger relevant signaling pathways for polarization and organization of the cytoskeleton [30]. Formation of such focal adhesion points is promoted within the silk network, where the sizes of the fibrils recapitulate the dimensions of ECM fibers with structures in the micrometer range [25]. Using combined staining of cells integrated in silk, we observe punctate vinculin-rich focal adhesions sequestered to the tips of cell protrusions (Fig. 3d, left, Suppl. Fig. 6), confirming integrin-involved binding of cells to the silk. In the hydrogel on the other hand, an integrin pair can bind to the RGD-motif, but the very thin alginate chains do not physically allow gathering of several integrins to the same spot, which prohibits formation of focal adhesions.

**Fig. 6.**
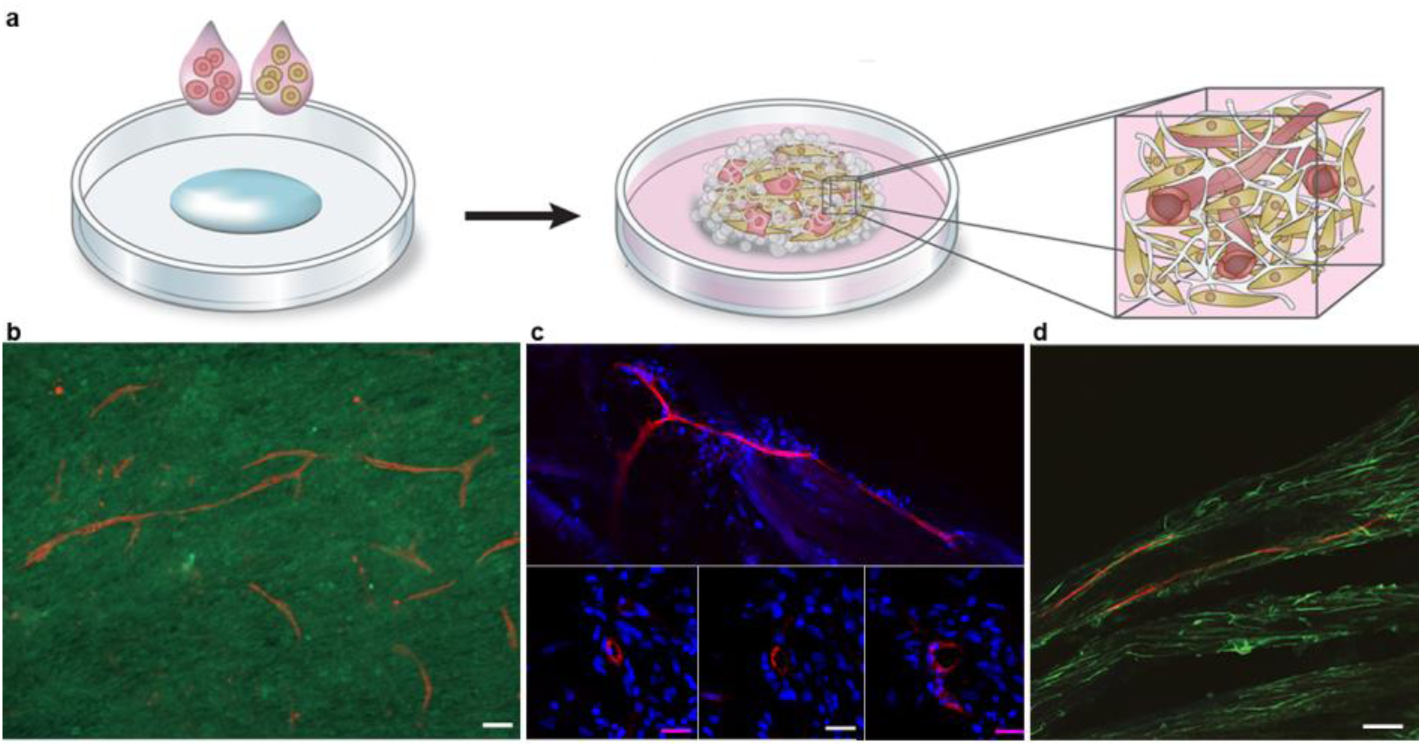
Formation of micro vessels within 3D silk. **(a)** The silk-assembly allows facile combination of two or more cell types. The schematics show an example where addition of a small fraction of endothelial cells together with a connective tissue cell type allows for vascularization of the resulting tissue construct. **(b)** Representative micrograph showing formation of long and branched vessel-like structures in FN-silk foam after 10 days co-culture of 2% endothelial cells (HDMEC, CD31, red) with mesenchymal stem cells (HMSC, CD44, green) in presence of isolated human pancreatic islets (not shown in the image). (N=5, n=2). Scale bar = 100 μm. **(c)** Incorporation of a fraction (10%) of endothelial cells (HDMEC) together with dermal fibroblasts (HDF) during formation of FN-silk fibers resulted in rearrangement into vessel-like structures during 35 days of culture (upper). Scale bar = 200 μm. Consecutive cryosections stained for CD31 (red) and nuclei (blue), shows that the vessel-like ring-formation appears at the same position in all three sections (lower). Scale bar = 25 μm (N=1-8, n=1-2). **(d)** Aligned CD31+ (red) cells were found in FN-silk fibers with endothelial cells (10%) and skeletal muscle cells after 14 days of culture.

## Silk fibers with integrated cells are highly extendable

The stiffness of the surrounding provides crucial signals that affect the fate of cells. The mechanical properties of biological tissue are complex, and especially the wide range of stiffness moduli, covering Pa in brain tissue [31], kPa in vessels [32], MPa in tendons [33] and GPa in cortical bone [34], is challenging to mimic *in vitro*. When compared with the stiffness of cell culture plates (1 GPa), it is not surprising that 2D culture on such substrates alter the properties of most cell types. Recently, it has been shown that tissue is remodulated by cell forces pulling the ECM fibers [25], showing the importance of flexible fibers within a scaffold to which cells can attach.

In order to obtain 3D cellular constructs, it is also of importance that the scaffold withstands attachment forces, media convections and handling by the personnel. The silk scaffolds with integrated cells constructed herein appeared strong and were stable enough for handling throughout the culture period and analysis procedures. In order to relate the mechanical properties to native tissue, silk fibers with integrated cells were subjected to tensile testing in a pre-warmed physiological buffer (Fig. 4a). Mesenchymal stem cells, known to attain an elongated cell shape and moderate production of exogenous ECM [35], were used as a model cell type for these tests. The fibers exhibited a linear plastic phase (Youngs modulus 1.8 +/-0.5 MPa) with up to 25% elongation. This was followed by a plastic phase with signs of irreversible deformations (Yield point 338 +/-107 kPa) and the silk fibers were extended to more than twice its initial length (strain 172 +/-73 %) before breakage (ultimate stress 450 +/-93 kPa) (Fig. 4b, Suppl. Fig. 7). These results suggest that the mechanical properties of silk with integrated cells match those of connective tissue [36], with most similarity to artery walls.

**Figure 7.**
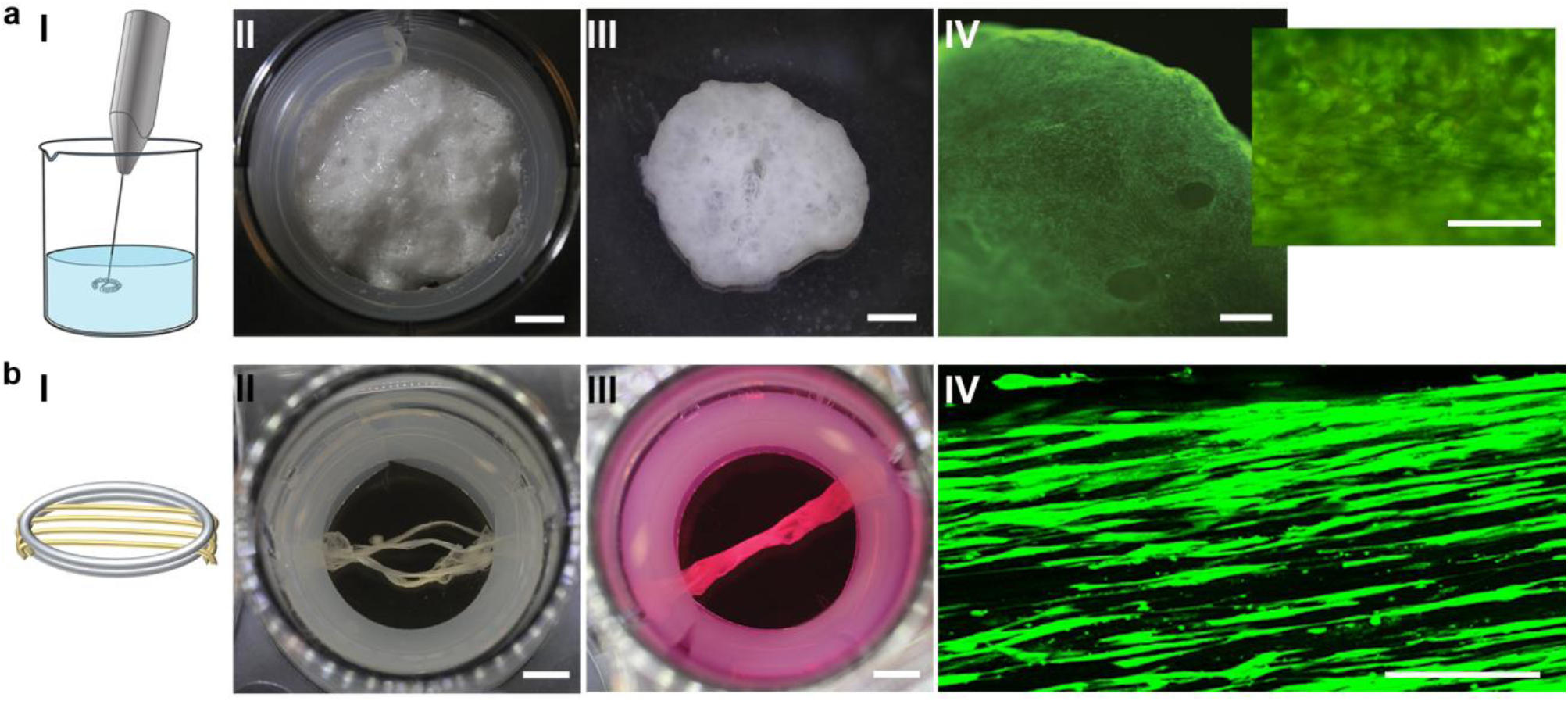
Silk-assembly with cells allows creation of macro-sized tissue constructs. **(a)** Larger foams can be obtained by using a small whisker to introduce air bubbles into the silk within a few seconds (I). A FN-silk foam with integrated HDF placed in a 6-well plate directly after whipping (II). Scale bar = 5 mm. After a few days the foam is free-floating in the well (III). Scale bar = 2.4 mm. Live stain shows a high level of viable cells after 14 days of culture (III). Scale bar = 600 μm. **(b)** Thicker fibers can be obtained by merging several fibers mounted in parallel (I). Several FN-silk fibers with integrated human muscle satellite cells (HSkMSC) and human endothelial cells were mounted in parallel in a 12 well plate (II), whereby they fuse within 7 days of culture (III). Scale bar = 4 mm. Confocal scan image of the middle plane of a fiber with integrated HDF at day 14 (lower), showing viable and aligned cells (green) inside of the fiber (IV). Scale bar = 50 μm.

Photographs taken during tensile testing confirms that the fibers are highly extendable (Fig. 4c) and that some filaments are ruptured before the whole bundle is broken. Micrographs of fiber pieces fixed and stained after tensile testing confirm elongated morphology of the integrated cells (Fig. 4d). Residuals of cells which seem to have their cytoskeleton snapped off can be seen along the ruptured fiber ends. This indicates that a force transition into and throughout the cells has occurred during stretching of the fiber, confirming proper cell attachment.

## Differentiation capacity of cells within 3D silk

Recent advances in methods for isolation and expansion of stem cells and the identification of inductive factors that direct differentiation along specific lineages has opened up the potential to provide an unlimited supply of cells for drug testing, research and medical transplantation. In order to engineer complete tissue models from stem cells, it is essential to create realistic artificial stem cell niches. The differentiation niches of stems cells *in vivo* are inherently 3D, and their biochemistry and topology strongly affect the differentiation process [37]. Therefore, we investigated the applicability of the herein described 3D culture set up for efficient differentiation, using both pluripotent and multipotent stem cells (Fig. 5).

Although more sensitive than most adult cell types, human embryonic stem cells could be viably integrated within the 3D silk foam. After 48 hours, expanding cell aggregates were found distributed throughout the 3D foam (Fig. 5b, left). Endodermal differentiation was then initiated, giving rise to dense layers of cells positive for the endodermal progenitor markers forkhead box protein A2 (FOX2A) and SOX17 (Fig. 5b, middle). Endodermal induction was further verified by expression analysis, confirming robust upregulation of *SOX17* and *CER1*, and down regulation of pluripotency (NANOG) (Fig. 5b, middle and right).

As example of multipotent cells, bone marrow-derived human mesenchymal stem cells (hMSC) were integrated within the 3D silk. After expansion, the cells were subjected to two protocols commonly used to steer the fate into adipogenic and osteogenic lineages, respectively (Fig. 5c). Lipid droplets were then found throughout silk scaffolds with integrated cells that had been treated with adipocyte induction media (Fig. 5c, left, Suppl. Fig. 8). Calcium phosphate was found deposited throughout cultures treated with osteoblast induction media, with accumulation throughout the fibers (Fig. 5c, right, Suppl. Fig. 8), confirming successful differentiation. In order to investigate the potential of 3D culture in silk for differentiation of adult precursor cells, human skeletal muscle satellite cells (HSkMSC) were expanded within silk fibers. After 2 weeks of further culture in myogenic differentiation media, the fibers were completely filled with aligned cells with prominent actin filaments (Fig. 5d, left). Elongated cells fused together with multiple nuclei and prominent expression of the muscle-specific marker desmin indicates myotube maturation (Fig. 5d, right).

## Co-culture with endothelial cells in 3D silk results in vessel formations

As 3D cultures grow in size, the access to nutrients and oxygen becomes increasingly hampered since the exchange is based on passive diffusion. In endogenous tissue, this supply is assured through the vasculature network. The lack of vessels thus limits 3D cultures to length scales under which oxygen gradients can occur [38]. The herein described method is practically convenient for direct combinations of many cell types, since this is easily achieved by the addition of several cell types to the silk protein solution (Fig. 6a). We examined what inherent organization capacity endothelial cells have for forming microvessels in a co-culture within silk. Before assembly, a fraction (2-10%) of endothelial cells was added together with cells of the connective tissue types (Fig. 6, Suppl. Fig. 9). Already within two weeks, endothelial cells had gathered and millimeter long, branched sprouts were found throughout a silk foam with co-cultured mesenchymal stem cells (Fig. 6b). Vessel-like structures with prominent rings of endothelial cells were also formed when co-cultured with dermal fibroblasts in silk fibers (Fig. 6c). Lumen formations (10-20 μm in diameter) resembling capillaries could be detected at the corresponding location in consecutive cryosections. Various states of vessel formations were also found aligned within the silk fibers after co-culture of endothelial cells and skeletal muscle cells (Fig. 6d).

## Silk assembly with cells allows creation of macro-sized 3D cultures

The so far presented experiments describe the potential of silk-assembly for 3D culture of cells in culture plates, and show applicability for research purposes. In order to demonstrate scalability of the methodology to broaden the application range, we will in the following section describe how larger constructs of silk with integrated cells can be achieved.

In order to obtain centimeter-sized, free-floating silk foams, we used a small whisker to introduce sufficient amount of tiny air bubbles within a few seconds (Fig. 7a, I). In this case, a suspension of fibroblasts in culture media was added just before whipping. For more sensitive cells, it is also possible to blend in the cells directly after whipping. By the use of a spoon, the resulting foam was placed in a culture well fitted with a silicone tube, to avoid silk attachment to the walls (Fig. 7a, II). After stabilization in the cell incubator for 1 h, the foam was covered with media. Within a few days, the foam detached to end up like a free-floating entity (Fig. 7a, III). After 14 days of culture, viable cells were found throughout the macro-sized foam (Fig. 7a, IV).

The length of the silk fibers obtained is directly dependent on the size of the lab trough used during formation, and thus easily scalable. However, the thickness of the fibers can only reach a certain level, limited by the concentration of protein and cells included. During culture of cells with contractile capacity, such as skeletal muscle cells, we noticed that the fibers curled up if kept free floating, confirming a cooperative action of the aligned cells. Therefore, we investigated the option to mount several fibers in parallel (Fig. 7b, I, II). Within a few days of culture, the fibers with integrated cells had fused to a single millimeter thick fiber (Fig. 7b, III). Confocal microscopy revealed a substantial amount of living cells throughout the depth of the fibers (Fig. 7b, IV).

## Outlook

We present a new strategy for integration of viable cells into 3D networks that mimics the fibrous architecture of native ECM. The mediator for this is the self-assembly of a silk protein that is recombinantly produced, and thereby defined and free from human and animal components. We herein report results showing that the silk formation method can be applied to a wide range of anchorage-dependent cell types, including endothelial cells, skeletal muscle cells, fibroblasts, keratinocytes as well as pluripotent and multipotent stem cells. The mild and accommodative silk-assembly process occurs simultaneous with cell integration, thereby allowing the cells to migrate and connect to each other. At the same time, a mechanically stable 3D support is provided by the silk. Thereby, intrinsic properties of native tissue, including proliferation in 3D, differentiation capacity and formation of vasculature networks, are reconstituted.

Cells gathered within 3D silk foam or cell sheets constitute a promising option for creation of glandular-like structures, while the fiber shape is more suitable for reconstruction of aligned structures found in blood/lymph vessels, nerves, muscle, ligament, tendon *etc.* The set-up and protocol for formulation of 3D silk with integrated cells is very simple and can be used by anyone without any need for special instrumentation or technical expertise. Moreover, it is convenient to perform the procedure under sterile conditions, within a laminar flow hood. Compared to time consuming techniques such as 3D-printing, the hands-on requirements for silk-assembly to integrate cells into macroscopic 3D scaffolds are easily fulfilled. The method is simple enough to be widely implemented as standard procedure in any cell lab, rendering 3D as easy to obtain as 2D cell culture.

## Methods

Full details are given in the Supplementary Information.

### 1 Recombinant spider silk protein

The spider silk protein FN-4RepCT kindly provided by Spiber Technologies AB was recombinantly produced in *E. coli* and purified using chromatographic methods.

### 2 Formulation of silk scaffolds with integrated cells

#### 2.1. Foam formation

FN-silk protein (3 mg/mL) were placed as a drop (15-40 μL) in the middle of a well of a non-treated, hydrophobic 24-well plate (Sarstedt). Air was quickly pipetted into the drop 20-30 times. Cells (0.5-2 ×10^6^ /mL) suspended in the appropriate culture media (containing 25mM Hepes, without serum), were added dropwise (10-20 μL) either before or after introduction of air bubbles. The foams were kept 15-60 minutes in the cell incubator before covered with complete cell culture medium. For foams integrated with hESC, 15 μl of FN-silk was used for 50 000 hESCs (15 000 cells/μl). Cells were integrated in the presence of 10 μg/ml Rock inhibitor Y27632 (VWR) and silk was stabilized for 15 min at 37°C in 5% CO2. After stabilization, 0.7 ml NutriStem supplemented with 10 μg/ml Rock inhibitor Y27632 was added. Rock inhibitor was omitted in the medium from 24 hours after seeding. Larger foams were prepared from 1.5 mL of FN-silk (3 mg/mL) subjected to 5-10 sec of whipping using a small battery operated hand whisker (Rubicson).

#### 2.2. Fiber formation

FN-silk protein (0.5-3 mg) was carefully mixed with cells (0.5-2 ×10^6^) suspended in the appropriate culture medium (containing 25mM Hepes, without serum) into a total volume of 2-4 mL. Fiber formation was induced by gentle repeated tilting of the lab trough for 1-3 hours at room temperature, while protected from direct light. The formed fibers were washed in pre-warmed 1xPBS and cut into 2-3 cm long pieces. The pieces were placed either mounted across or free-floating in non-treated 12 or 24-well plate with fresh medium during further culture. In order to obtain thicker fibers, several pieces (4-8) were mounted next to each other, enabling fusion during culture.

#### 2.3. Sheet formation

Silk sheets were made by adding 5-20 μL FN-silk protein (3 mg/mL) in the middle of a well of a non-treated, hydrophobic 24-well plate (Sarstedt). An equal volume of cells suspended in appropriate culture medium (0.5-1 ×10^6^/mL) was added directly to the drop of protein. The sheets were left for 30-60 min in the cell incubator, followed by 30 min without lid in the laminar flow hood, before 1 mL of complete culture medium was added for further culture.

### 3. Mechanical analysis

Uni-axial tensile test measuring force versus strain (% length extension) of fibers with integrated cells (n=13) was performed under physiological-like conditions (37+/-1°C, 1xPBS) using a servo-electric custom built Zwick/Roell material testing machine equipped with a 50 N class 1 load cell. The tests were performed on fibers with integrated MMSCs after culture for 14-24 days. Once the 1xPBS buffer bath was heated up, the load cell was leveled and the fiber ends gently mounted in stainless-steel grippers aligned with the machine load line. The tests were performed at a fixed displacement rate of 0.01 mm/s, corresponding to a strain rate of about 0.1%/s. Displacement and load were continuously recorded until the specimen fractured, *i.e.* the load dropped. From the recorded tensile load *F*, the First Piola-Kirchhoff stress *P=F/A* in kPa was calculated, where the fiber’s initial cross-sectional area, *A*, was calculated from fiber dimensions measured with a light microscope (Suppl. Table 3). The strain average (ε) was extracted from the total deformation of the fibers measured at the grip close to the fiber end using video extensometers. The strain *ε=100·u/L 0* was calculated by relating the grip displacement u to the fibers initial test length *L0* Fibers that broke at the clamps were not considered for the analysis of mechanical properties.

## Acknowledgements

This work was supported by Knut & Alice Wallenberg foundation, The Swedish Research Council, VINNOVA, Formas and The Novo Nordisk foundation. Spiber Technologies AB is acknowledged for providing soluble silk proteins. Selina Parvin is acknowledged for experimental assistance and Prof. P-Å Nygren for valuable comments on the manuscript.

## Author contributions

M.H. conceived and directed the research. M.H., U.J. and M.W. designed the experiments. U.J., M.W., N.D.S., C.P.T. and C.Å. performed cell experiments and analyses. C.G. and I.L.A. designed and supervised the mechanical testing. L.N. designed the illustrations. M.H., M.W. and U.J. wrote the manuscript. All authors discussed the results and commented on the manuscript.

## References

1. Lee, J., M.J. Cuddihy, and N.A. Kotov, Three-dimensional cell culture matrices: state of the art. Tissue Eng Part B Rev, 2008. 14(1): p. 61–86.

2. Ghosh, S., et al., Three-dimensional culture of melanoma cells profoundly affects gene expression profile: a high density oligonucleotide array study. J Cell Physiol, 2005. 204(2): p. 522–31.

3. Kenny, P.A., et al., The morphologies of breast cancer cell lines in three-dimensional assays correlate with their profiles of gene expression. Mol Oncol, 2007. 1(1): p. 84–96.

4. Mazzoleni, G., D. Di Lorenzo, and N. Steimberg, Modelling tissues in 3D: the next future of pharmaco-toxicology and food research? Genes Nutr, 2009. 4(1): p. 13–22.

5. Weigelt, B., et al., HER2 signaling pathway activation and response of breast cancer cells to HER2-targeting agents is dependent strongly on the 3D microenvironment. Breast Cancer Res Treat, 2010. 122(1): p. 35–43.

6. Broutier, L., et al., Human primary liver cancer-derived organoid cultures for disease modeling and drug screening. Nat Med, 2017. 23(12): p. 1424–1435.

7. Ravi, M., et al., 3D cell culture systems: advantages and applications. J Cell Physiol, 2015. 230(1): p. 16–26.

8. Onoe, H. and S. Takeuchi, Cell-laden microfibers for bottom-up tissue engineering. Drug Discov Today, 2015. 20(2): p. 236–46.

9. Elbert, D.L., Bottom-up tissue engineering. Curr Opin Biotechnol, 2011. 22(5): p. 674–80.

10. Morimoto, Y., A.Y. Hsiao, and S. Takeuchi, Point-, line-, and plane-shaped cellular constructs for 3D tissue assembly. Adv Drug Deliv Rev, 2015. 95: p. 29–39.

11. Zorlutuna, P., et al., Microfabricated biomaterials for engineering 3D tissues. Adv Mater, 2012. 24(14): p. 1782–804.

12. Billiet, T., et al., A review of trends and limitations in hydrogel-rapid prototyping for tissue engineering. Biomaterials, 2012. 33(26): p. 6020–41.

13. Rowley, J.A., G. Madlambayan, and D.J. Mooney, Alginate hydrogels as synthetic extracellular matrix materials. Biomaterials, 1999. 20(1): p. 45–53.

14. Baker, B.M. and C.S. Chen, Deconstructing the third dimension - how 3D culture microenvironments alter cellular cues. Journal of Cell Science, 2012. 125(13): p. 3015–3024.

15. Stark, M., et al., Macroscopic fibers self-assembled from recombinant miniature spider silk proteins. Biomacromolecules, 2007. 8(5): p. 1695–1701.

16. Hedhammar, M., et al., Sterilized recombinant spider silk fibers of low pyrogenicity. Biomacromolecules, 2010. 11(4): p. 953–9.

17. Fredriksson, C., et al., Tissue Response to Subcutaneously Implanted Recombinant Spider Silk: An in Vivo Study. Materials, 2009. 2(4): p. 1908–1922.

18. Widhe, M., N.D. Shalaly, and M. Hedhammar, A fibronectin mimetic motif improves integrin mediated cell biding to recombinant spider silk matrices. Biomaterials, 2016. 74: p. 256–266.

19. Widhe, M., et al., Recombinant spider silk as matrices for cell culture. Biomaterials, 2010. 31(36): p. 9575–85.

20. Nilebäck L, A.S., Kvick M, Paananen A, Linder MB, Hedhammar M Interfacial behavior of recombinant spider silk protein parts reveals cues on the silk assembly mechanism. Submitted manuscript, 2018.

21. Hedhammar, M., et al., Structural properties of recombinant nonrepetitive and repetitive parts of major ampullate spidroin 1 from Euprosthenops australis: implications for fiber formation. Biochemistry, 2008. 47(11): p. 3407–17.

22. Ushiki, T., Collagen fibers, reticular fibers and elastic fibers. A comprehensive understanding from a morphological viewpoint. Archives of Histology and Cytology, 2002. 65(2): p. 109–126.

23. Kierszenbaum, A., Histology and cell biology-an introduction to pathology. 2nd ed. 2002: Mosby.

24. Askarieh, G., et al., Self-assembly of spider silk proteins is controlled by a pH-sensitive relay. Nature, 2010. 465(7295): p. 236–8.

25. Baker, B.M., et al., Cell-mediated fibre recruitment drives extracellular matrix mechanosensing in engineered fibrillar microenvironments. Nat Mater, 2015. 14(12): p. 1262–8.

26. McGuigan, A.P. and M.V. Sefton, Vascularized organoid engineered by modular assembly enables blood perfusion. Proc Natl Acad Sci U S A, 2006. 103(31): p. 11461–6.

27. Bohari, S.P., D.W. Hukins, and L.M. Grover, Effect of calcium alginate concentration on viability and proliferation of encapsulated fibroblasts. Biomed Mater Eng, 2011. 21(3): p. 159–70.

28. Khattak, S.F., et al., Application of colorimetric assays to assess viability, growth and metabolism of hydrogel-encapsulated cells. Biotechnol Lett, 2006. 28(17): p. 1361–70.

29. Guillame-Gentil, O., et al., Engineering the extracellular environment: Strategies for building 2D and 3D cellular structures. Adv Mater, 2010. 22(48): p. 5443–62.

30. Khetan, S., et al., Degradation-mediated cellular traction directs stem cell fate in covalently crosslinked three-dimensional hydrogels. Nat Mater, 2013. 12(5): p. 458–65.

31. Rashid, B., M. Destrade, and M.D. Gilchrist, Mechanical characterization of brain tissue in compression at dynamic strain rates. J Mech Behav Biomed Mater, 2012. 10: p. 23–38.

32. Forsell, C., et al., Biomechanical properties of the thoracic aneurysmal wall: differences between bicuspid aortic valve and tricuspid aortic valve patients. Ann Thorac Surg, 2014. 98(1): p. 65–71.

33. Wren, T.A., et al., Mechanical properties of the human achilles tendon. Clin Biomech (Bristol, Avon), 2001. 16(3): p. 245–51.

34. Spatz, H.C., E.J. O’Leary, and J.F. Vincent, Young’s moduli and shear moduli in cortical bone. Proc Biol Sci, 1996. 263(1368): p. 287–94.

35. Amable, P.R., et al., Protein synthesis and secretion in human mesenchymal cells derived from bone marrow, adipose tissue and Wharton’s jelly. Stem Cell Research & Therapy, 2014. 5.

36. Gohl, K.L., A. Listrat, and D. Bechet, Hierarchical mechanics of connective tissues: integrating insights from nano to macroscopic studies. J Biomed Nanotechnol, 2014. 10(10): p. 2464–507.

37. Fuchs, E., T. Tumbar, and G. Guasch, Socializing with the neighbors: stem cells and their niche. Cell, 2004. 116(6): p. 769–78.

38. Rodenhizer, D., et al., A three-dimensional engineered tumour for spatial snapshot analysis of cell metabolism and phenotype in hypoxic gradients. Nat Mater, 2016. 15(2): p. 227–34.

